# Hesperetin inhibits foam cell formation in macrophages via activating LXRα signal in an AMPK dependent manner

**DOI:** 10.1101/2020.01.22.915819

**Authors:** Xuanjing Chen, Dezhi Zou, Xiaoling Chen, Huanlin Wu, Danping Xu

## Abstract

Cholesterol efflux from macrophages is the first step of cholesterol reverse transport (RCT), whose increase inhibits cholesterol accumulation and foam cell formation to suppress atherogenesis. Liver X receptor alpha (LXRα) and adenosine monophosphate activated protein kinases (AMPK) both have the pivotal role in cholesterol homeostasis. However the association between these two molecules in cell model of atherosclerosis is poorly understood. Hesperetin has been reported to possess several protective effects for cardiovascular diseases, while little is known about the role of hesperetin and its underlying mechanism on macrophage foam cell formation. In this study, we sought to investigate the potential effects of hesperetin in cholesterol efflux by using human macrophage derived foam cells, focusing on liver X receptor alpha (LXRα) and adenosine monophosphate activated protein kinases (AMPK) implication. Hesperetin treatment concentration-dependently reduced foam cell formation, intracellular cholesterol level and cholesterol esterification rate, and enhanced cholesterol efflux in THP-1 macrophages. Hesperetin upregulated the protein levels of LXRα and its targets including ABCA1, ABCG1 as well as SR-BI, and phosphorylated-AMPK. Meanwhile, hesperetin-induced upregulation of LXRα expression was enhanced by AMPK agonist and inhibited by AMPK inhibitor. Furthermore, hesperetin increased mRNA level of LXRα and its target genes, all which were depressed by AMPKα1/α2 small interfering RNA (siRNA) transfection. In conclusion, we founded for the first time that hesperetin could active AMPK. And this activation upregulated LXRα and its targets including ABCA1, ABCG1 and SR-BI, which significantly inhibited foam cell formation and promoted cholesterol efflux in THP-1 macrophages. Our results highlight the therapeutic potential of hespretin for the possible reduction in foam cell formation. This new mechanism could contribute the anti-atherogenic effects of hesperetin.

## Introduction

Reverse cholesterol transport (RCT) is known as an effective approach to alleviate hypercholesterolemia and atherosclerosis (1). It refers to the process by which excess cholesterol from lipid laden peripheral cells is transferred to plasma high-density lipoprotein and then transporting to liver for bile acid synthesis and is finally excreted via the feces (2).

Liver X receptors (LXRs) is ligand-activated nuclear transcription factor containing two isoforms, LXRα and LXRβ. LXRα predominates in the tissues and cells related to lipid metabolism, such as liver, intestine, adipose tissue and macrophages (3). LXRα has been demonstrated to play a pivotal role in transcriptional regulation of cholesterol homeostasis by regulating the transcription of genes that encode proteins involved in cholesterol biosynthesis, catabolism, secretion as well as reverse transport (4). Cellular cholesterol efflux, the initial step of RCT, is primarily mediated by three identified target proteins of LXRα namely ATP-binding cassette subfamily A member 1 (ABCA1), ATP-binding cassette transporters ATP-binding cassette subfamily G member 1 (ABCG1) and scavenger receptor class B type I (SR-BI) (5, 6). Activation of LXRα signal pathway by endogenous lipid ligands or synthetic agonist like T0901317 promotes cellular cholesterol efflux from lipid-laden macrophages to attenuate atherosclerosis progression (7, 8).

LXRs transcriptional activity, like other nuclear receptors, is regulated by post-translational modifications including acetylation, SUMOylation, and phosphorylation (9–10). Previous work has linked LXRs phosphorylation to some kinases, such as mitogen-activated protein kinases, protein kinases C, casein kinases, and adenosine monophosphate activated protein kinases (AMPK) (11–13). AMPK is a ubiquitously expressed serine/threonine protein kinase complex responsible for maintaining balance between anabolic and catabolic pathways for cellular energy homeostasis (14). Once activated, AMPK triggers the catalytic progress to genatate ATP, which control the biosynthesis that consume ATP (15). Considerable evidences indicated that AMPK activation inhibited the formation of foam cells and the deposition of atherosclerotic plaque (16, 17). However, the association between AMPK and LXRα in cell model of atherosclerosis is still to be illustrated.

Flavonoids compounds are known to have multiple biological and pharmacological activities (18). Hesperetin, a citrus bioflavonoid, is a major bioactive component from Fructus Aurantii and Pericarpium Citri Reticulatae that both are medical herb using in China for thousands of years to against heart diseases and digestive disease (19–21). It has been shown that hesperetin have prominent role in anti-inflammation, anti-oxidation, anti-apoptosis, insulin-sensitizing as well as lipid-lowering, and exhibit a protective effect on cardiovascular system, nervous system as well as metabolic homeostasis of lipid and glucose (22–26). Several evidences showed that hesperetin could reduce cholesterol accumulation and enhance RCT (27–29). However, the underlying mechanism is still largely obscure. Recently, hesperetin has been suggested as the potent bioactivator of LXRα and AMPK (30, 31). Based on these facts, we hypothesised that LXRα and AMPK signal pathway may be involved in the effect of hesperetin on RCT. Therefore, we examined the effect of hesperetin on foam cell formation, cholesterol level and cholesterol efflux in THP-1 macrophages. Furthermore, we investigated the underlying mechanisms that the transcription and translation level of LXRα as well as its downstream targets, and the implication of AMPK in this process.

## Materials and methods

### Materials

RPMI-1640 medium, fetal bovine serum(FBS), and BCA assay kit were purchased from Thermo Fisher Scientific (Waltham, MA, USA). Phorbol 12-myristate 13-acetate (PMA), T0901317 (293754-55-9), Compound C (866405-64-3), AICA-Riboside (3031-94-5), 3-(4,5-Dimethyl-2-thiazolyl)-2,5-diphenyl-2H-tetrazolium bromide (MTT) (298-93-1), Oli Red O (1320-06-5), and phenylmethanesulfonyl fluoride (PMSF) were purchased from Sigma-Aldrich (St. Louis, MO,USA). Cholesterol Efflux Assay Kit (ab196985), anti-LXR alpha antibody (ab176323), anti-ABCA1 antibody (ab18180), anti-ABCG1 antibody (ab52617), and anti-Scanvenging Receptor SR-BI antibody (ab217318) were purchased from Abcam (Cambridge, MA, USA). RIPA lysis buffer, anti-Phospho-AMPKα antibody(50081), anti-GADPH antibody(5174), anti-rabbit IgG, HRP-linked antibody (7074), and anti-mouse IgG HRP-linked antibody (7076) were purchased from Cell Signaling Technology (Danvers, MA, USA). Polyvinylidine difluoride membrane and ECL kit (WBKLS0500) were purchased from Merck Millipore (Billerica, MA, USA). Control siRNA (sc-37007), AMPK alpha 1 siRNA (sc-29673), and AMPK alpha 2 siRNA (sc-38923) were purchased from Santa Cruz Biotechnology (Dallas, TX, USA). Total RNA Extraction Reagent (R401-01), HiScript® II Q RT Super Mix (Q711-02/03), and ChamQ Universal SYBR qPCR Master Mix (Q711-02) were purchased from Vazyme Biotech Ltd. (Nanjing, China). Tissue total cholesterol assay kit (E1015) and tissue free cholesterol assay kit (E1016) were purchased from Applygen Technologier Inc. (Beijing, China). Oxidized low dentisty lipoprotein (ox-LDL), hesperetin (520-33-2), phosphatase inhibitor cocktail, and anti-AMPK alpha antibody (66536-1-lg) were purchased from Yiyuan Biotechnologies (Guangdong, China), Chengdu Chroma-Biotechnology Ltd. (Sichuan, China), Roche (Basel, Switzerland), and Proteintech Group Inc. (Rosemont, USA) respectively.

### Cell culture and treatments

THP-1 human monocytic leukemia cell line was obtained from Cell Bank of the Chinese Academy of Sciences (Shanghai, China). THP-1 cells were grown in RPMI-1640 supplemented with 10% FBS, 100 U/mL penicillin, 100 μg/mL streptomycin, 2.5 μg/mL amphotericin B and 2 mM L-glutamine under a humidified atmosphere at 37 °C and 5% CO2. To facilitate differentiation into macrophages, cells (1×10^6^ cells/mL) were cultured with 160nM phorbol12-myristate 13-acetate for 24h. THP-1-derived macrophages were then incubated with 80μg/mL ox-LDL combined with different treatments for another 24h.

Hesperetin, T0901317, AICAR and compound C, were prepared in a minimal volume of dimethyl sulfoxide (DMSO) and stored in refrigerator at -20℃. For treatment, reagents were added into cultured medium and final concentration of DMSO was less than 0.1%.

### Cell viability

THP-1 cells were seeded in 96-well plate at a density of 1 × 10^4^ per well and cultured overnight. The cells were then treated with gradient dilutions of hesperetin (0-1000 μM) for 24h. 5mg/mL of MTT in RPMI-1640 medium was added to each well of plate and incubated with cells at 37°C for 4h. Subsequently, the medium was discarded and incubation with DMSO for 10min at room temperature. The absorbance was measured at 570nm with a microplate reader (BioTek Instruments Inc, Winooski, USA).

### Oil red O staining and lipid accumulation analysis

THP-1 cells were seeded in 6-well plates at a density of 2 × 10^5^ per well with treated as described above. Subsequently, THP-1 derived macrophage foam cells were rinsed 3 times with cold phosphate-buffered saline (PBS) and then fixed in 4% paraformaldehyde for 30min. After rinsed 3 times with distilled H2O, the cells were stained with 0.5 mL fresh filtered Oil Red O solution (0.5% Oil red O in 60% isopropanol) per well for 30min at room temperature. Foam cells were observed and photographed by light microscopy (OLYMPUS BX53, Olympus, Tokyo, Japan).

After Oil Red O staining, intracellular stained-lipid droplets were extracted by isopropanol. The absorbance was measured at 492 nm with a microplate reader (BioTek Instruments Inc, Winooski, USA). The quantitative results were corrected after parallel experiments of cellular protein contents.

### Analysis of cellular cholesterol accumulation and cholesterol efflux

The level of cellular TC and FC were determined using Tissue total cholesterol assay kit and Tissue free cholesterol assay kit. All experiments were performed following the manufacturer’s instructions. The contents of cholesteryl ester (CE) were determined by subtracting FC from total cholesterol. Cellular protein contents were calculated according to a BCA assay kit. Macrophage cholesterol efflux capacity was determined using a Cholesterol Efflux Assay Kit. The procedure was according to the manufacturer’s instructions. Brifely, THP-1 macrophages were pretreated with fluorescently-labeled cholesterol for 6h and then treated with or without hesperetin for 24h. Cell supernatant was collected and cells were solubilized with cell lysis buffer. Microplate reader was used to analyzed the absorbance of the supernatant and lysates at 490 nm. The percentage of cholesterol efflux is the ratio between fluorescence intensity of media and fluorescence intensity of cell lysate plus media × 100%.

### Western blot analysis

Cells in 6-well plates were lysed with RIPA lysis buffer containing 1mM PMSF and 1% phosphatase inhibitor cocktail (PhosSTOP, Roche group, Swiss). The supernatant was collected after centrifuging at 14,000 rpm for 15 min at 4℃. The concentration of cellular supernatant was examined by Pierce BCA Protein Assay Kit. Equal amounts of proteins (60ug each) were separated on 10% sodium-dodecyl sulfate polyacrylamide gel electrophoresis (SDS-PAGE) and transferred to polyvinylidine difluoride membrane at 300 mA for 1.5 h at 4℃. Membranes were incubated with primary antibodies overnight at 4℃ after 1h blocking in 5% skimmed milk (Becton, Dickinson and Company, USA) at room temperature. Membranes were then washed with TBST for total 15 min and incubated with horseradish peroxidase (HRP) conjugated second antibodies for 1h at room temperature. Protein bands were detected by enhanced chemiluminescence method using ECL kit, and band intensity was analysed by Image Lab 5. 2. 1 software (Bio-Rad, Hercules, USA).

Primary antibodies include: anti-AMPK alpha antibody, anti-LXR alpha antibody, anti-ABCA1 antibody, anti-ABCG1 antibody, anti-Scanvenging Receptor SR-BI antibody, anti-GADPH antibody. Second antibodies include: anti-rabbit IgG HRP-linked antibody, anti-mouse IgG HRP-linked antibody.

### Transfection of small interfering RNA

THP-1 macrophages were transfected with non-silencing control siRNA or siRNA targeting AMPKα1/α2 according to the manufacturer’s instructions. Briefly, THP-1 derived macrophages were seeded into 6-well plates at a density of 2 × 10^5^ cells/well. Added the siRNA duplex solution to the dilute transfection regent, and incubated the transfection reagent mixture 30 minutes at room temperature. After washing the cells once with transfection medium, cells were incubated with transfection reagent mixture in transfection medium (siRNA final concentration at 60nM) at 37℃ for 6h. And then 1mL of RPMI-1640 medium containing 2 times the normal serum and antibiotics concentration was added to each wells and incubated with the cells for another 24h. Subsequently, cells were treated with different reagents for 24h. Target genes knockdown was validated by real time RT-PCR, using GADPH expression vector as an internal control.

### Real-time RT-PCR analysis

THP-1 macrophages were seeded in 6-well plates. After transfection and treatments as described above, total mRNA was extracted using Total RNA Extraction Reagent. First-strand cDNAs were synthesized from 1ug of total RNA using a HiScript® II Q RT Super Mix. The 2 μL of cDNAs were amplified by ChamQ Universal SYBR qPCR Master Mix in a total volume of 25 μL. Real-time quantitative polymerase chain reaction was carried out by applying a BioRed CFX96 with the specific thermocycler condition following the manufacturer’s instructions. GADPH was used as the internal control and quantitative measurements were analyzed using the ΔΔCt method. The primers sequences are presented in Table. 1.

**Table 1.**
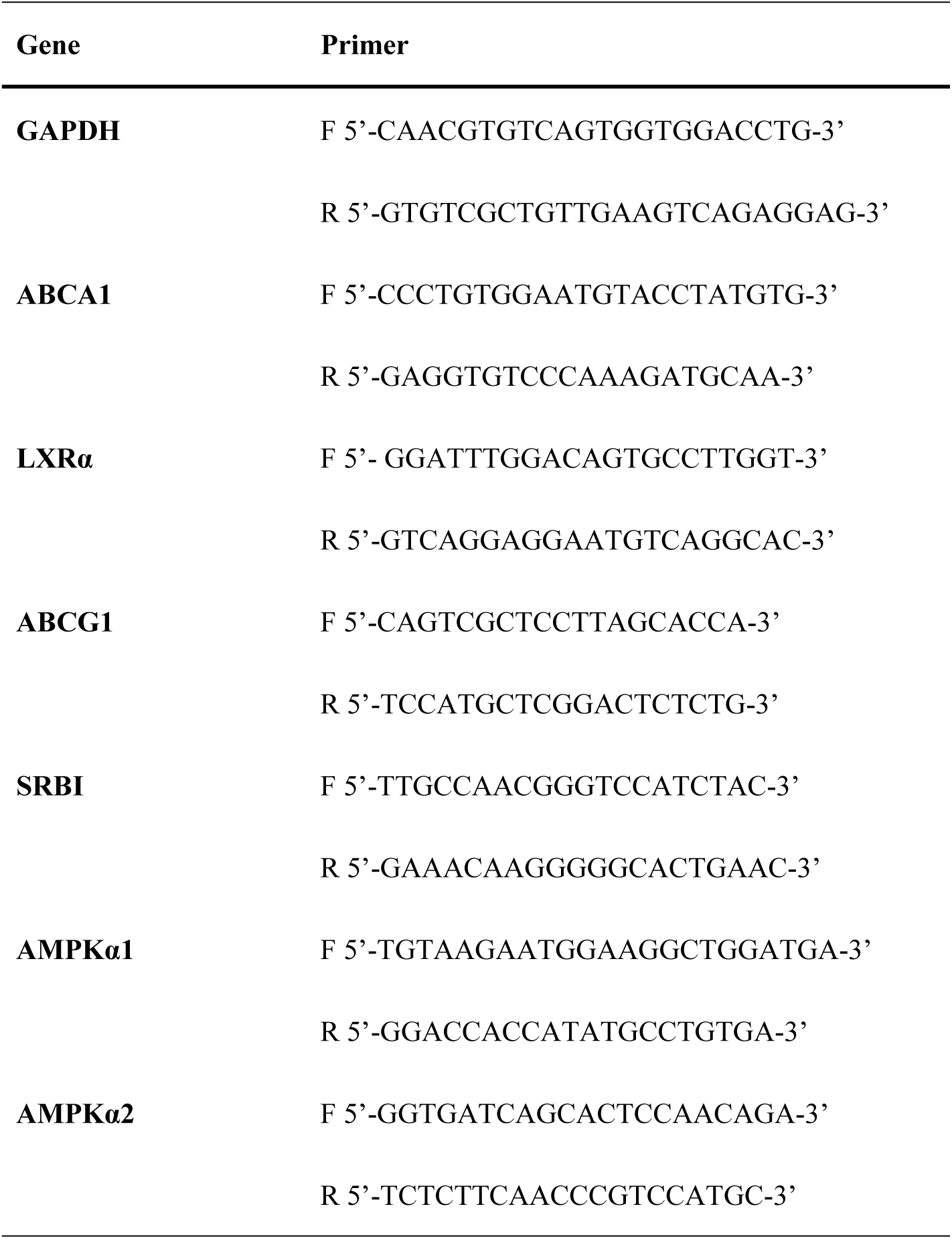
Sequences of primer for real-time RT-PCR

### Statistical analysis

All data are presented as the mean ± standard deviation (SD) from at least three independent experiments. The statistical significance of differences between groups was analyzed using one-way ANOVA and student’s t-test analysist by SPSS version 19.0 software. Differences with a *P* value less than 0.05 were considered statistically significant.

## Results

### Hesperetin inhibits foam cell formation and reduces lipid accumulation in THP-1 macrophages

Firstly, we observed the effects of hesperetin in different concentrations (0-1000 μM) on THP-1 macrophage activity. Data showed that high concentration of hesperetin could reduce the viability of THP-1 macrophages, and a maximum hesperetin concentration of 100uM had no effects on cell viability (Figure 1A). So we chose 10 μM, 50 μM and 100 μM concentrations of hesperetin to carry out next experiments.

**Figure 1.**
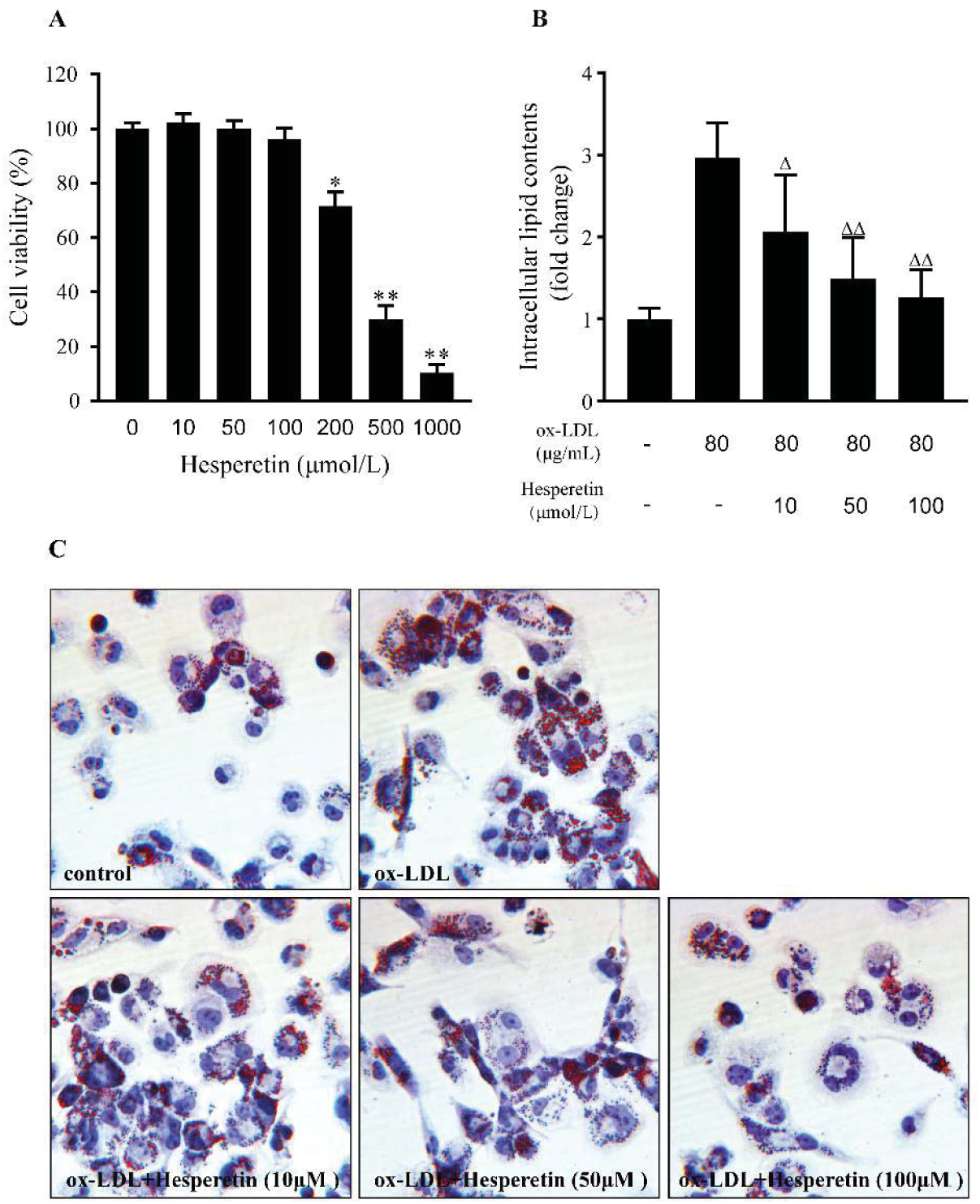
Effect of different hesperetin concentrations on cell viability in THP-1 macrophages. (**A**) Cell were treated with gradient dilutions of hesperetin (0-1000μmol/L) and subject to cell viability experiment by MTT method. Concentration-dependent effect of hesperetin on lipid accumulation in THP-1 macrophage foam cells. THP-1 macrophages were treated with 80 μg/mL ox-LDL alone or combined with hesperetin (10μM, 50 μM and 100 μM) for 24h. Control cells were cultured for the same time without any additions. (**B**) Representative microscopy image of Oil Red O-stained lipid droplets were observed at a magnification of × 200. (**C**) Semi-quantification of intracellular lipid contents were determined by the absorbance of intracellular pigments extracted by isopropanol at 492nm. Results are expressed as mean ± SD from five independence experiments performed in triplicate. \**p* < 0.05, ** *p* < 0.01 vs. control; Δ *p* < 0.05, ΔΔ *p* < 0.01 vs. treatment with 80 μg/mL ox-LDL alone.

To investigate the effect of hesperetin on foam cell formation and lipid accumulation, 80μg/mL ox-LDL with or without hesperetin were incubated with THP-1 macrophages for 24h. And then the Oil Red O-stained lipid droplets in cells were observed. Data showed that hesperetin significantly reduced the size and amount of lipid droplets in THP-1 macrophage foam cells in a dose dependent way (Figure 1C). Quantitative analysis of intracellular lipid contents is determined by isopropanol extraction of intracellular pigment. Result showed that hesperetin dose-dependently reduced lipid contents in THP-1 macrophage foam cells (Figure 1B).

### Hesperetin reduces cholesterol accumulation and promots cholesterol efflux in THP-1 macrophage foam cells

Since hesperetin exhibited the significant effect on reducing lipid accumulation. We supposed that hesperetin would also inhibit cholesterol accumulation. As shown in Figure 2A and Figure 2B, compared with control cells, total cholesterol (TC), cholesterol ester (CE) as well as esterification rate (CE/TC) were significantly increased after ox-LDL incubation. However, hesperetin dose-dependently reduced intracellular TC, CE as well as esterification rate.

**Figure 2.**
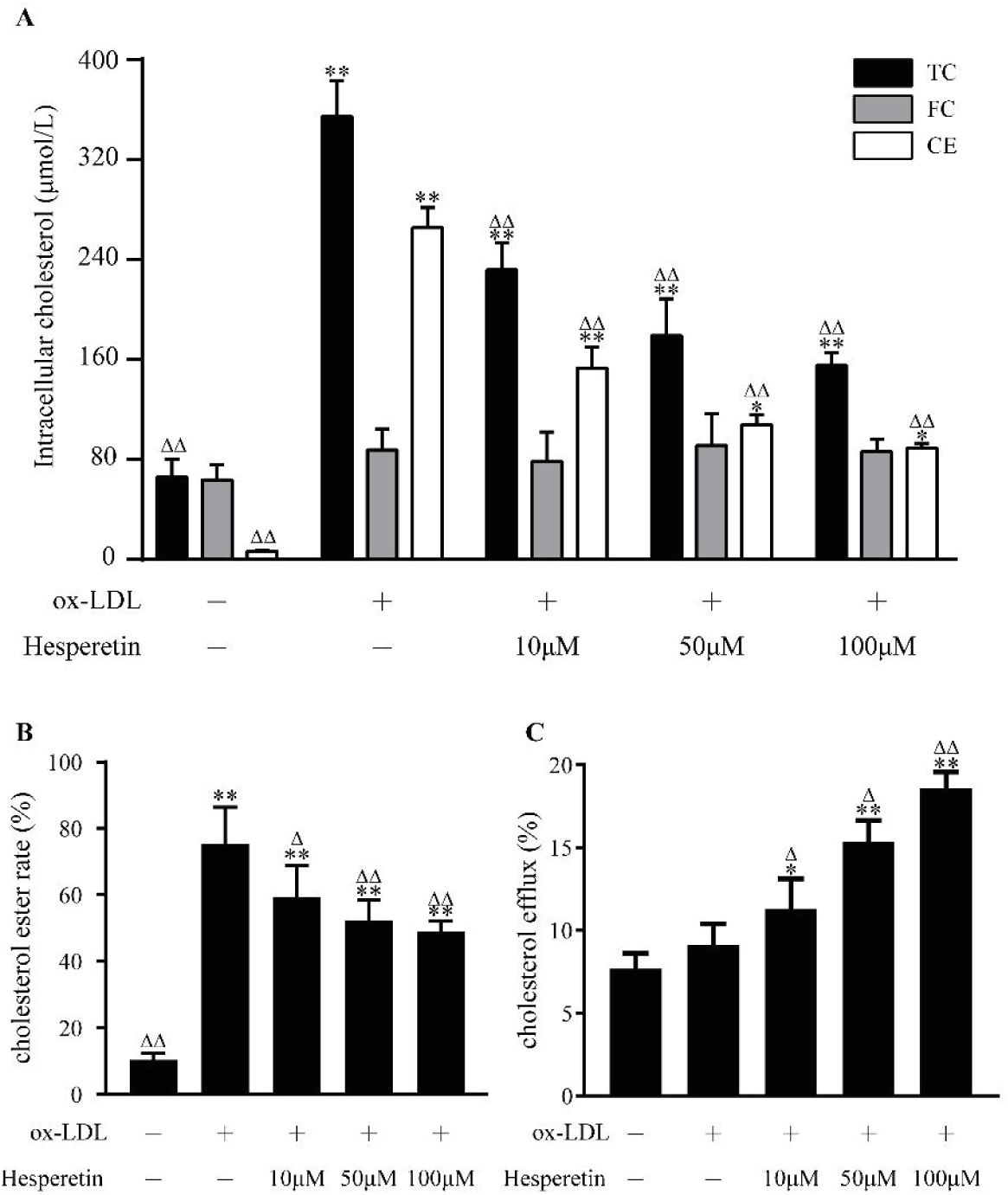
Concentration-dependent effect of hesperetin on cholesterol accumulation and cholesterol efflux in THP-1 macrophage foam cells. THP-1 macrophages were treated with 80 μg/mL ox-LDL alone or combined with hesperetin (10 μM, 50 μM and 100 μM) for 24h. Control cells were cultured for the same time without any additions. (**A**) The levels of total cholesterol (TC), free cholesterol (FC) and cholesterol ester (CE) in THP-1 macrophage foam cells were examined by enzymic method; (**B**) Cholesterol esterification rate (CER) is the ratio of CE to TC; (**C**) The commercial assay was performed to determine cholesterol efflux as described above. Results are expressed as mean ± SD from five independence experiments with each performed in triplicate. \**p* < 0.05, ** *p* < 0.01 vs. control; Δ *p* < 0.05, ΔΔ *p* < 0.01 vs. treatment with 80 μg/mL ox-LDL alone.

Furthermore, we assessed cholesterol efflux ability to investigate how hesperetin reduces intracellular cholesterol level. As shown in Figure 2C, hesperetin increased cholesterol efflux to 11.31%, 15.38% and 18.58% at concentrations of 10 μM, 50 μM, 100 μM respectively, all which are significantly different from cells with ox-LDL alone and control group.

### Hesperetin up-regulates the protein and mRNA expression of LXRα in THP-1 macrophages

LXRa signal is demonstrated to have critical role in cellular cholesterol homeostasis. So we observed the influence of hesperetin on LXRα protein and mRNA level. As shown in Figure 3A and 3B, hesperetin dose-dependently increased LXRα protein and mRNA expression while this increase was lower than T0901317. We chose the best effective concentration of hesperetin at 100uM to further validate its role in activating LXRα. Interestingly, when cells were treated with hesperetin and T0901317, a synthetic effect on LXRα was produced both in protein and mRNA level, which has a significantly difference to treatment with T0901317 alone.

**Figure 3.**
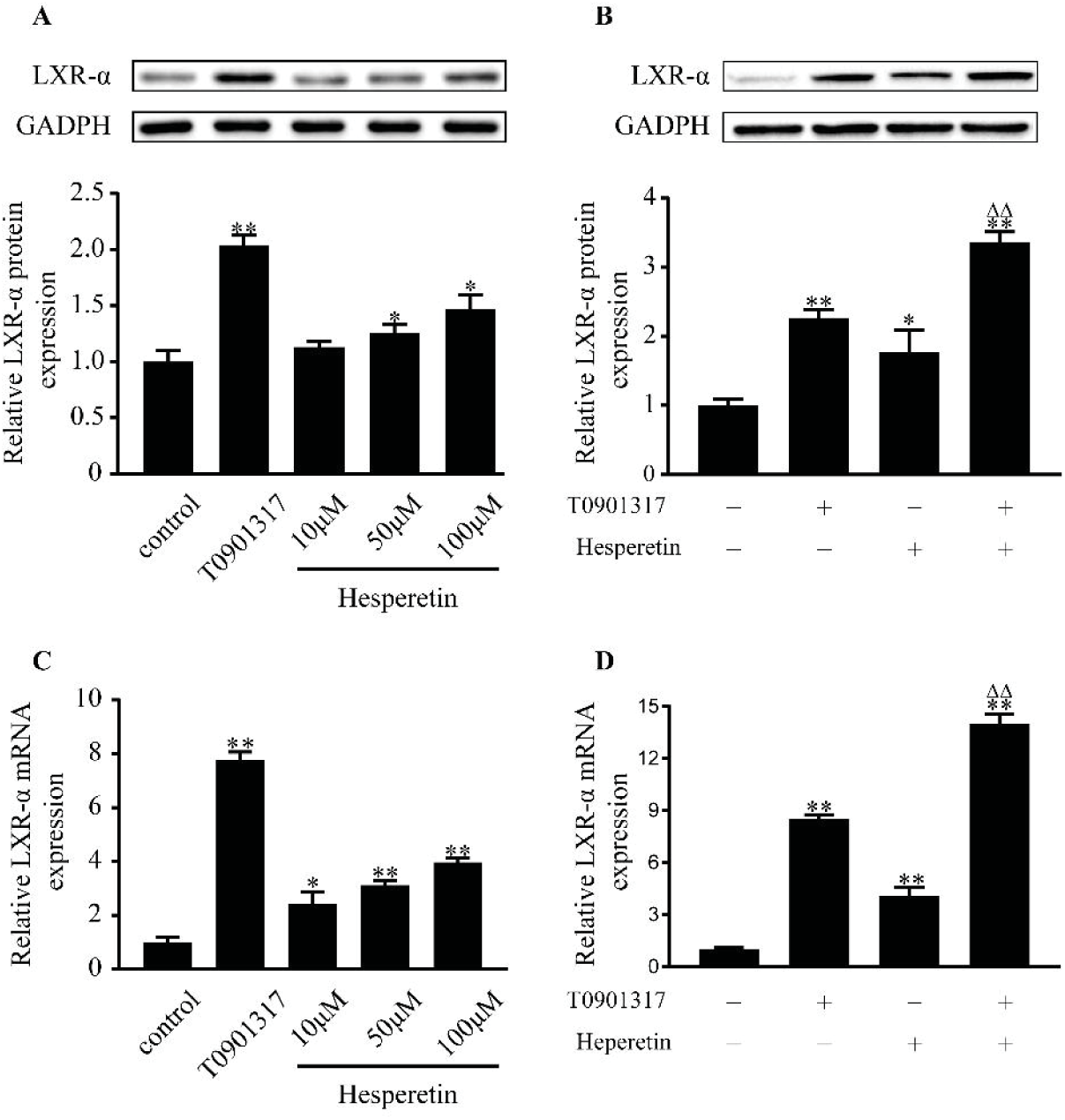
Effect of hesperetin on LXRα expression in THP-1 macrophages. THP-1 macrophages were treated with hesperetin (10 μM, 50 μM, 100 μM) and/or T0901319 (10 μM) for 24h. Control cells were cultured for the same time without any additions. Total proteins and RNAs were extracted and analyzed by western blot and real-time RT-PCR. (A and C) Concentration-dependent effect of hesperetin on LXRα protein and mRNA expression; (B and D)The best effctive concentration of hesperetin at 100 μM was used in further validating hesperetin-induced activation of LXRα. Results are expressed as mean ± SD from three independence experiments with each performed in triplicate. \**p* < 0.05, ** *p* < 0.01 vs. control; Δ *p* < 0.05, ΔΔ *p* < 0.01 vs. treatment with T0901317.

### Hesperetin influences the expression of LXRα target proteins in THP-1 macrophages

We further examined the role of hesperetin in LXRα downstream target proteins including ABCA1, ABCG1, and SR-BI. Data showed that hesperetin dose-dependently increased ABCA1, ABCG1 and SR-BI protein level. Among these, the up-regulation of ABCG1 by hesperetin was slightly higher than by T0901317. However, the up-regulation of ABCA1 and SR-BI were lower than by T0901317 (Figure 4A and 4B).

**Figure 4.**
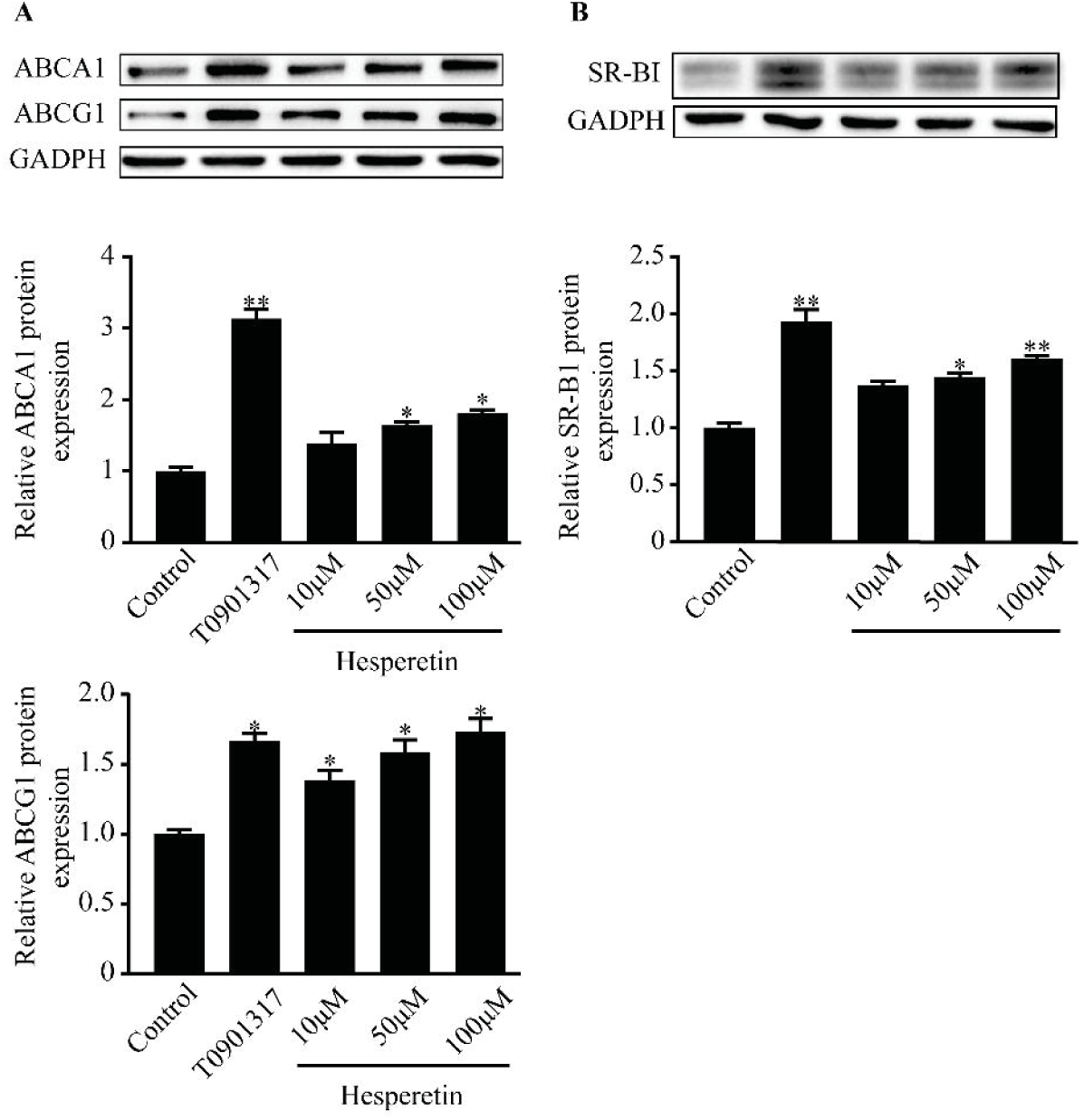
Effect of hesperetin on LXRα target proteins. THP-1 macrophages were treated with hesperetin (10 μM, 50 μM, 100 μM) or T0901317(10 μM) for 24h. Control cells were cultured for the same time without any additions. Total proteins were extracted and subject to western blot analysis. (**A**) Concentration-dependent effect of hespretin on the protein expression of ABCA1 and ABCG1; (**B**) Concentration-dependent effect of hespretin on the protein expression of SR-BI. Results are expressed as mean ± SD from three independence experiments with each performed in triplicate. * *p* < 0.05, ** *p* < 0.01 vs. control; Δ *p* < 0.05, ΔΔ *p* < 0.01 vs. treatment with T0901317.

### Hesperetin actives AMPK in THP-1 macrophages

AMPK has been described as a key regulator of lipid synthesis. Therefore the effect of hesperetin on AMPK activation was examined. As shown in Figure 5A, hesperetin siginificantly upregulates phosphorylated-AMPK(pAMPK) protein level by 1.5 times compared with control group in THP-1 macrophages. Meanwhile a synthetic effect is observed when AICAR combined with hesperetin. Under this treatment, pAMPK protein expression is increased by 2.5 times compared with control and have significant difference to treatment with AICAR alone. However, there are no statistic difference in total AMPK protein level between groups (Figure 5B).

**Figure 5.**
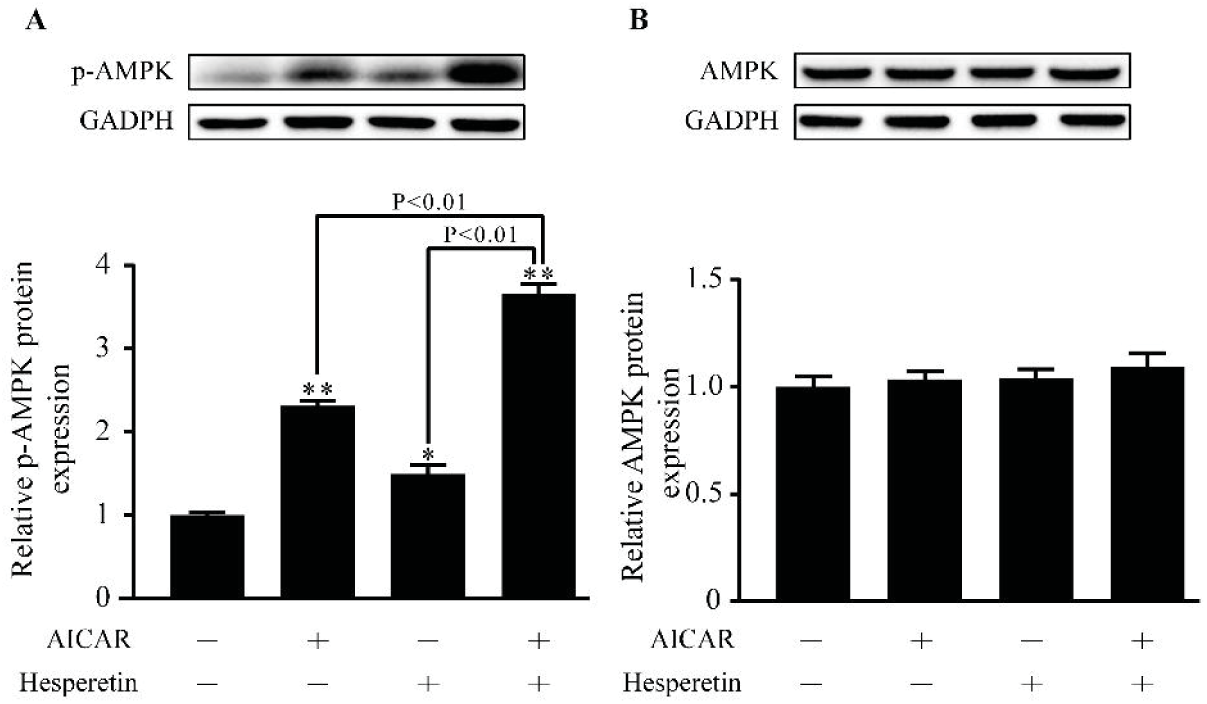
Effect of hesperetin on AMPK activation. THP-1 macrophages were treated with hesperetin (100 μM) and/or AICA-Riboside (AICAR) (100 μM). Control cells were cultured for the same time without any additions. (**A**) Phosphorylated-AMPK and (**B**)AMPK protein expression were analyzed by western blot. Results are expressed as mean ± SD from three independence experiments with each performed in triplicate. \**p* < 0.05, ** *p* < 0.01 vs. control.

### AMPK is implicated in LXRα activation induced by hesepretin

Previous study has demonstrated that LXRα may be regulated by phosphorylation of AMPK. We therefore investegated whether the upregulaion of AMPK induced by hesperetin has an effective role in LXRα activation. The agonist (AICAR) and inhibitor(Compound C) were performed. Results showed that AICAR could increase protein expression of LXRα induced by T0901317 or hesperetin (Figure 6A). While Compound C decreased protein expression of LXRα induced by T0901317 or hesperetin (Figure 6B). Similar situation was observed in mRNA level (Figure 6C and 6D).

**Figure 6.**
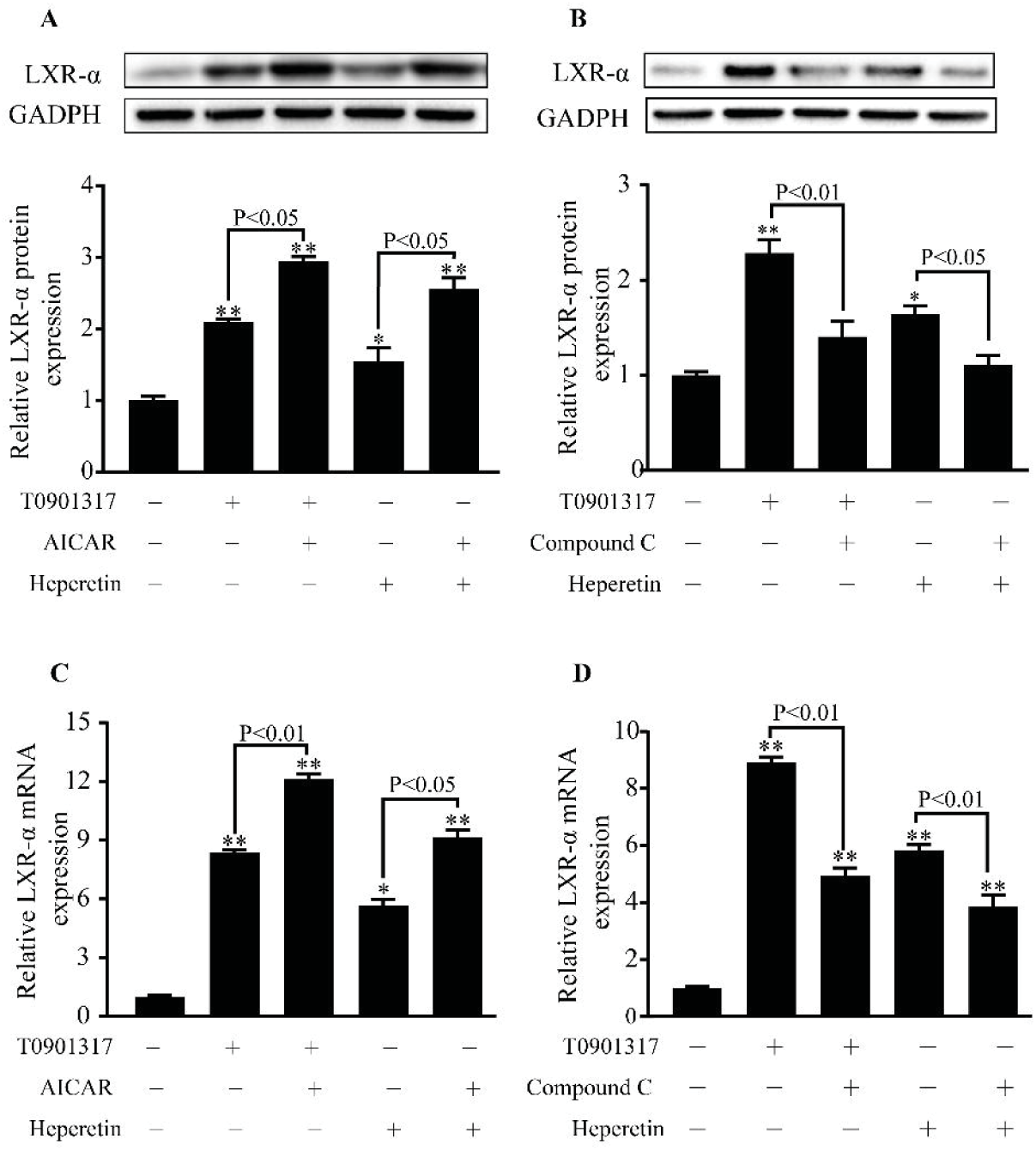
The role of AMPK in LXRα activation induced by hesperetin in THP-1 macrophages. THP-1 macrophages were treated hesperetin (100 μM) with or without AICARA or Compound C. Control cells were cultured for the same time without any additions. (**A** and **C**) The effect of AMPK activation on the up-regulation of LXRα protein and mRNA level induced by T0901317 or hespretin. (**B** and **D**) The effect of AMPK inhibition on the up-regulation of LXRα protein and mRNA level induced by T0901317 or hespretin. Results are expressed as mean ± SD from three independence experiments with each performed in triplicate. \**p* < 0.05, ** *p* < 0.01 vs. control.

### Transfection with AMPKα1/α2 siRNA inhibits the activation of LXRα signal induced by hesperetin

THP-1 macrophages were transfected with specific AMPKα1/α2 siRNA to further validate the role of AMPK in hesperetin-induced the upregulation of LXRα and its target genes. Control siRNA that will not lead to specific degradation of mRNA expression was used as a control group. After transfection with AMPKα1/α2 siRNA, AMPKα1 and AMPKα2 mRNA expression were significantly decreased (Figure 7A). This transfection also reduced the mRNA expression of LXRα as well as its target genes including ABCA1, ABCG1 and SR-BI when compared with transfection of control siRNA. Similar changes were observed in cells treated with T0901317 or hesperetin. And a synthetic effecte on LXRα and its target genes were founded in treatment with T0901317 and hesperetin (Figure 7B-7E). Bsed on this results and previous evidences in protein level. We therefore concluded that hesperetin-induced activation of LXRα signal pathway could be related to its activation of AMPK.

**Figure 7.**
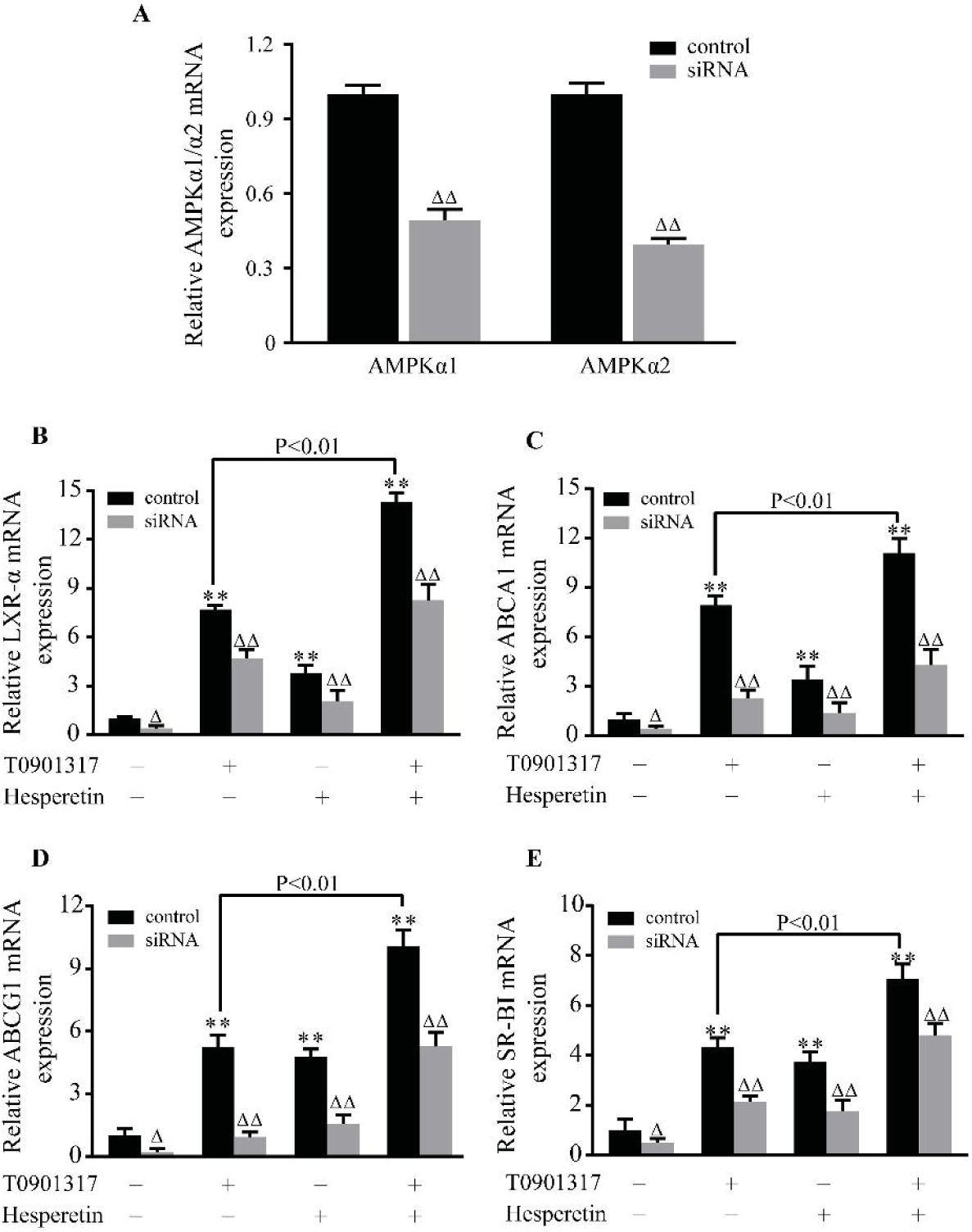
The role of AMPKα1/α2 siRNA in the regulation of LXRα and its target genes by hesperetin. THP-1 macrophages were transfected with control siRNA or siRNA against AMPKα1/α2. And then the cells were treated with hesperetin (100 μM) and/or T0901317 (10 μM). Control cells were cultured for the same time without any additions. (A-F) The mRNA expressions of AMPKα1, AMPKα2, LXRα, ABCA1, ABCG1, SR-BI were examined by real-time RT-PCR as described above. Results are expressed as mean ± SD from three independence experiments with each performed in triplicate. \**p* < 0.05, ** *p* < 0.01 vs. control; Δ *p* < 0.05, ΔΔ *p* < 0.01 AMPKα1/α2 siRNA vs. control siRNA.

## Discussion

In present study, we founded for the first time that hesperetin could activate AMPK in THP-1 macrophages. And this activation upregulated LXRα and its downstream targets including ABCA1, ABCG1 as well as SR-BI, which significantly inhibited foam cell formation, reduced lipid accumulation as well as cholesterol accumulation, and promoted cholesterol efflux in THP-1 macrophages.

The early stages of atherosclerosis is featured by lipoprotein cholesterol accumulation and foam cells formation within blood vessel walls (1). It has been reported that the incidence of cardiovascular events in patients of the Dallas Heart Study was inversely associated with cholesterol efflux (32). Numerous studies demonstrated that hesperetin exhibited evidently hypolipidemic, anti-inflammatory and endothelial protective properties (33, 34). Recently, Noriko Sugasawa et al. showed that hesperetin evidently attenuated the development of atherosclerotic lesions in apolipoprotein E knockout mice (35). Our results showed that hesperetin could enhance cholesterol efflux from THP-1 macrophages, and also reduced foam cells formation, cholesterol accumulation as well as esterification in THP-1 macrophages, which could contribute to the observed anti-atherogenic and hypolipidemic potential of hesperetin.

Numerous studies identified that LXRα was a promising anti-atherogenic target due to its roles in inhibiting inflammation and promoting reverse cholesterol transport (7). Activation of LXRα induces the expression of ABCA1, ABCG1 and SR-BI, all which are the membrane pumps exporting cholesterol to extracellular acceptors. Hesperetin has been reported to upregulate ABCA1 mRNA expression in THP-1 macrophages (30). Our study is consisted with this results and added the findings that hesperetin could significantly upregulate mRNA expression of LXRα and its target genes including ABCG1 as well as SR-BI. And all this upregulations were validated at protein level. Meanwhile we founded that unlike LXRα agonist-induced the preferential up-regulation of ABCA1 over ABCG1, hesperetin upregulated this two transporters in a similar level. This facts suggested that aside from LXRα signal there were trans-activating factors that specially control the transcriptional regulation of ABCG1 in these cells. Besides, it is worth noting the synthetic effect on LXRα signal when T0901317 is combined with hesperetin. Similar effects were demonstrated in other flavonoids such as the effect of quercetin on mRNA and protein levels of LXRα and its targets SRBI in HepG2 cells and C57BL/6 mice, and the upregulation of ABCG5 mRNA expression induced by genistein (36, 37).

Previous studies have demonstrated that phosphorytion of AMPK could up-regulate LXRα expression (38). Flavonoids, such as naringenin and quercetin, were considered as the effective therapeutic approaches that appear to influence AMPK (39, 40). Recently, Hajar Shokri Afra1 et al. founded that hesperetin could actives SIRT1-AMPK signaling pathway in HepG2 cells (31). In present study, we founded for the first time that hesperetin significantly up-regulated phosphorylated-AMPK protein expression. And addition of hesperetin to AICAR up-regulated phosphorylated-AMPK protein level by 1.5 times compared with treatment with AICAR alone. This results demonstrated a clear effect of hesperetin on AMPK activation in THP-1 macrophages. Based on the multiple bio-functions of AMPK signal, we suggested hesperetin might be a promising therapeutic strategy aimed at activation AMPK signal pathway in various pathological process.

We next examined the role of AMPK in hesperetin-induced LXRα activation. Hesperetin or T0901317 were combined with or without AMPK synthetic agonist (AICAR) or inhibitor (Compound C) to treat THP-1 macrophages. And then the changes of LXRα expression were observed. Results showed that hesperetin-induced LXRα activation could be enhanced by AICAR and inhibited by Compound C. Similar situation was observed in T0901317 treatment. Furthermore specific AMPKα1/α2 siRNA transfection was performed to validate the role of AMPK in LXRα and its target genes. Results showed that AMPKα1/α2 siRNA transfection significantly reduced hesperetin- and/or T0901317-induced mRNA upregulation of LXRα and its targets genes including ABCA1, ABCG1, and SR-BI. We therefore conclude that AMPK could be involved in the regulation of LXRα activation by hesperetin. Our results were in line with previous observations that AMPK activated LXRα in human macrophages (41, 42). However, there were several studies demonstrated that AMPK inhibited LXRα to prevent lipid accumulation in HepG2 cells (43, 44). This discrepancy may be due to the different characteristic properties between pathological cell models, which suggests AMPK may have a selective regulation on LXRα in different pathological process.

In conclusion, our study demonstrates that hesperetin concentration-dependently prevents foam cell formation, reduces total cholesterol level as well as cholesterol esterification rate, and enhances cholesterol efflux in THP-1 macrophages, which is due to the activation of LXRα signal pathway in an AMPK dependent manner. This new mechanism contributes to the beneficial role of hesperetin in regulating cholesterol homeostasis. We suppose that like other flavonoids, hesperetin could naturally find its place in the routine treatment of hyperlipemia and atherosclerosis.

## Acknowledgments

This study was funded by National Natural Science Foundation of China (No.81774219) and Guangzhou Science, Technology and Innovation Commission (No.).

## Abbreviations

ABCA1: ATP-binding cassette subfamily A member 1
ABCG1: ATP-binding cassette subfamily G member 1
ABCG5: ATP-binding cassette subfamily G member 5
AMPK: adenosine monophosphate activated protein kinases
CE: cholesterol ester
CER: Cholesterol esterification rate
LXRα: liver X receptor alpha
pAMPK: phosphorylated-AMPK
RCT: cholesterol reverse transport
SR-BI: scavenger receptor class B type I
TC: total cholesterol

